# Ancient genomes from Ladakh reveal 2800-year-old mixture between Tibetans and South Asians

**DOI:** 10.64898/2026.01.30.702804

**Authors:** Nick Patterson, Veena Mushrif-Tripathy, Quentin Devers, Lijun Qiu, Sonam Dolma, Gregory Soos, Swapan Mallick, Nadin Rohland, David Reich

**Author notes:** Corresponding author. (N.P.), (V. M.-T.), (Q.D.), (D.R.). Contributed equally (Nick Patterson, Veena Mushrif-Tripathy, Quentin Devers).

## Abstract

Reconstructing population history is harder in South Asia than in many other world regions due to a paucity of ancient DNA. We report genome-wide data for ten individuals from Old Lady Spider Cave, which lies 4000 meters above sea level in the Himalayan region of Ladakh, and dates to around 1500 years before present (BP). These individuals were genetically homogeneous and had an ancestry signature rare in South Asians today: admixed in roughly 50-50% proportions between a population well-proxied by present-day North Indians, and another genetically similar to ancient Tibetans. By analyzing the typical sizes of segments of DNA inherited from each of these ancestral populations, we find that mixture of these groups began at least fifty generation before the date of the individuals, that is, by around 2800 BP.

**Teaser:** Discovery of a previously unknown population that lived 2800-1500 years ago in the Himalayan region of Ladakh.

## Introduction

The Indian Union Territory of Ladakh lies at the crossroads of ancient trade routes connecting the Tibetan Plateau, parts of Central Asia including the Tarim Basin and the Hindukush-Pamir corridor, Kashmir, and South Asia. Petroglyphs dating to the Bronze Age (5000-3000 BP) document cultural links with Central Asian cultures, such as the Okunevo and Afanasievo of the Altai and south Siberia, and the Andronovo of west Central Asia (*1-5)*. Petroglyphs dating to the Iron Age (3000-2000 BP) in the “Animal” or “Steppic Style”, provide links with the Saka cultures of the west Central Asian steppes (*1, 3, 5*).

After 2300 BP, an east-west divide appears in the material culture record (*6, 7*). The east of Ladakh is characterized by “Corded Ware” style ceramics (not to be confused with the 5000-4400 BP European culture of the same name) found in the wider West-Tibetan and West-Himalayan world, including in Spiti, Guge, Purang, and Mustang. The west of Ladakh, where Old Lady Spider Cave is located, is marked by Central Asian-influenced ceramics with red slips and distinctive shapes and decoration, occurring in Hellenistic (2300-2000 BP), Kushan (2000-1600 BP) and post-Kushan (1600-1100 BP) times (*5*). There are also Kharoṣṭī and Brāhmī inscriptions, typical of the Kushan and post-Kushan world (*8*). According to oral traditions, Ladakh was once inhabited by speakers of language groups including Dardic (Indo-European speakers found today across northwest India, northern Pakistan, and northeast Afghanistan), Mon (across the Himalayan arch from India to Burma), Turkic, Mongol, and Tibetic. A limitation is that we do not know when any of the speakers of these languages first came to Ladakh, and these ethnonyms must reflect only a subset of the groups in the region in prehistoric times.

The Old Lady Spider Cave, or Abi Srinjamo Bao, is located at an altitude of 4000 meters in the pastureland above the village of Yogma Kharbu, and is one of the few deep caves of the region (Figure 1). A first large room contains petroglyphs and remains of looted cist graves. A long and narrow tunnel leads to a second large room, where bones are densely scattered in two main clusters. A third room can be accessed from the second room and its ceiling is only 50 cm high. Bones are also scattered in this room, although more sparsely. On the basis of highly preserved skeletal elements like hand and foot bones, it is likely that the bodies were kept on the cave floor and not moved after their primary deposition (*9*). As the cave has been visited by many people for centuries, however, the bones are disturbed. One of the main aims of the excavation was to verify the context of the bones and to determine whether bones were present below the existing floor of the cave (*9*).

**Figure 1:**
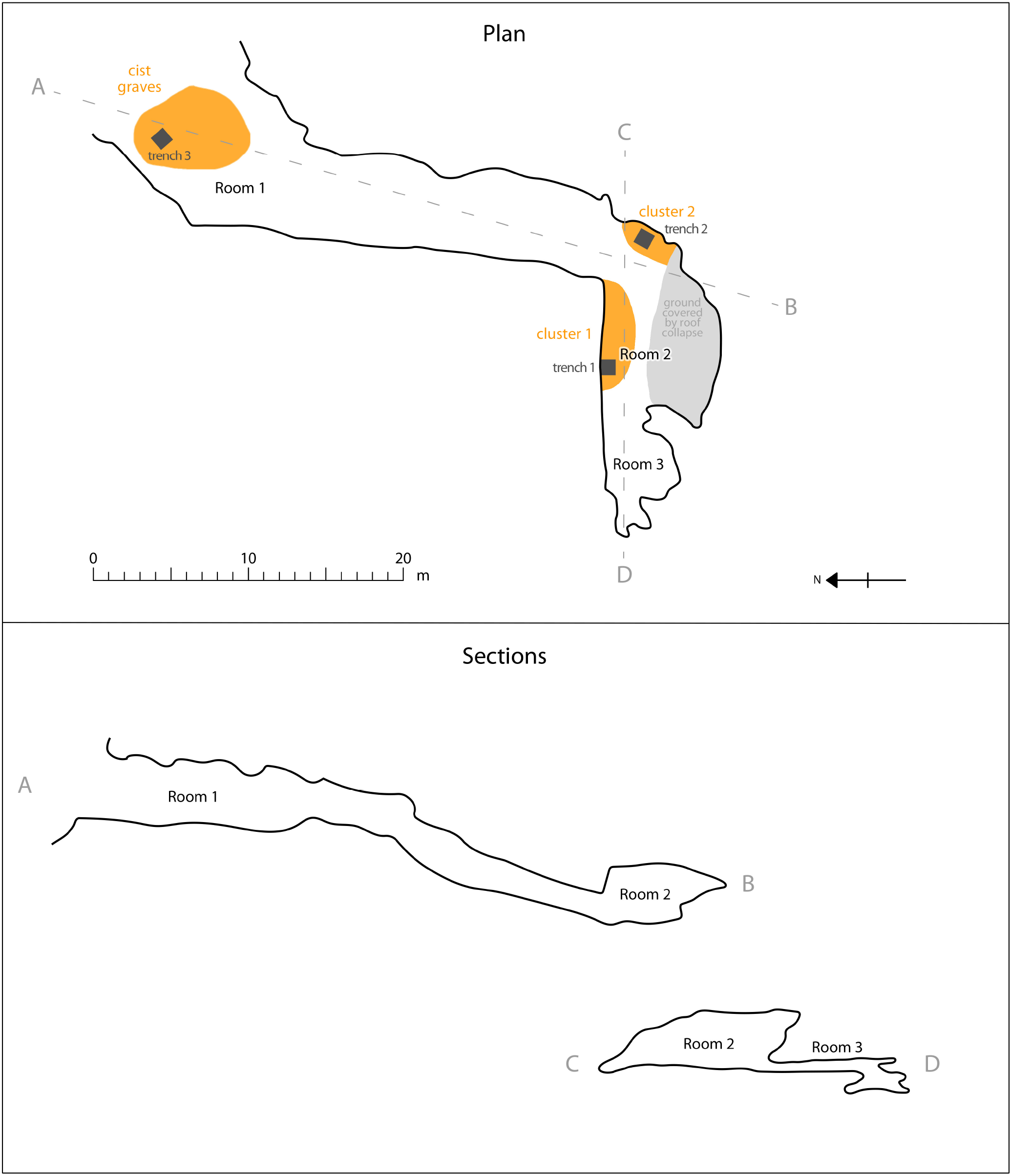
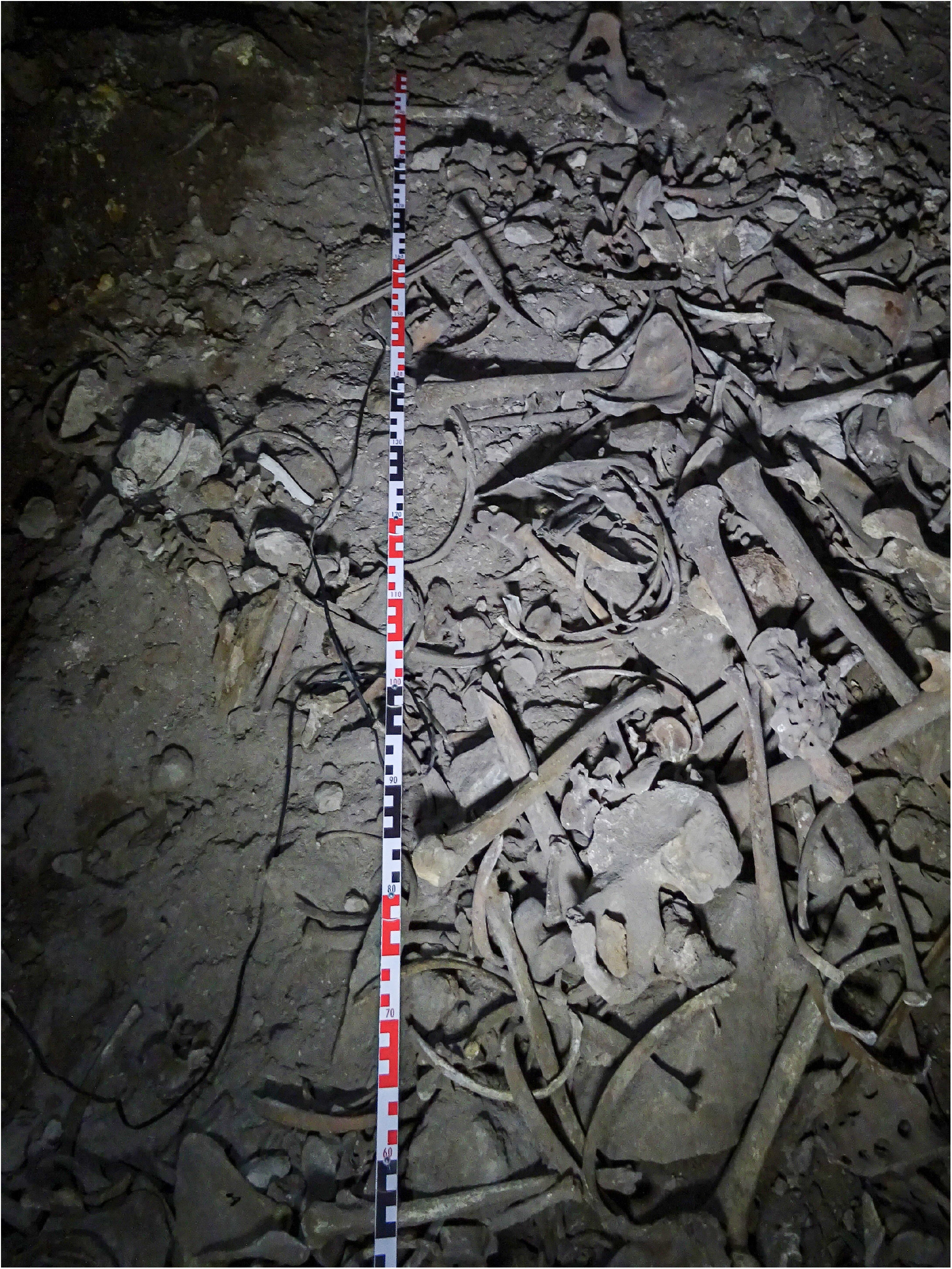
The Old Lady Spider Cave. (a) Plan and section of the cave, drawn from a previously unpublished point-cloud shared with permission by Danilo Rosati. (b) Previously unpublished photograph of bones in room 2, taken by the authors during the excavation.

Excavations were conducted in 2021 by a group directed by two of the authors (Veena Mushrif-Tripathy and Quentin Devers), under the auspices of the Archaeological Survey of India (ASI). The collection represents all age groups (infants to elders) and both adult males and females (*9*). Other findings include bones with naturally mummified tissues (skin, ligaments, muscles), including two mummified hands. With the goal of minimizing contamination, we wrapped bones and mummified tissues in clean aluminum foil and transferred them to clean plastic bags at the Deccan College Post-Graduate and Research Institute (Pune). We obtained permission from the ASI to export the samples to Harvard University in order to study them using specialized ancient DNA preparation and in-solution enrichment procedures. Author Veena Mushrif-Tripathy hand-carried the samples to Boston USA for this analysis.

### Data set assembly

We extracted DNA from 11 specimens using protocols adapted to the specific type of sample (bone or soft tissue). The samples were 8 bones (a clavicle, a femur, a fibula, a humerus, a radius, a tarsal, an ulna, and a rib), and 3 pieces of soft tissue (all from the pelvic area) (*10-13*). We generated 52 double-stranded (*14*) and 12 single-stranded (*15*) sequencing libraries, enriched them in solution using the “Twist Ancient DNA” assay targeting 1,352,535 single nucleotide polymorphisms (SNPs) (*16*), and sequenced them on Illumina NovaSeq instruments aiming for 20-30 million paired sequences of 101 base pairs each for each library (Table S1). We aligned the sequences to the human reference genome, and carried out almost all of our analyses at a set of 584,131 SNPs on chromosomes 1-22 that had also been genotyped in hundreds of relevant modern populations using the Affymetrix Human Origins SNP genotyping array (*17*).

After quality control and merging data obtained from two samples that we determined based on the genetic results to be from the same individual (I36588 and I36589, referred to in Table 1 as I36588), we had data from 10 unique individuals. Six represented by bones yielded excellent data with 0.85-12.5-fold average sequence coverage on targeted positions (median 6.3-fold). One individual represented by a pelvic tissue sample had usable but not particularly high-quality data (0.048-fold coverage). The remaining three individuals represented by two pelvic tissue samples and one clavicle had too little data for genome-wide analysis (0.0016-0.015-fold) (Table 1).

**Table 1:**
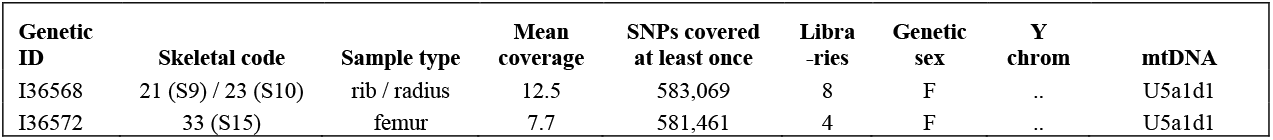
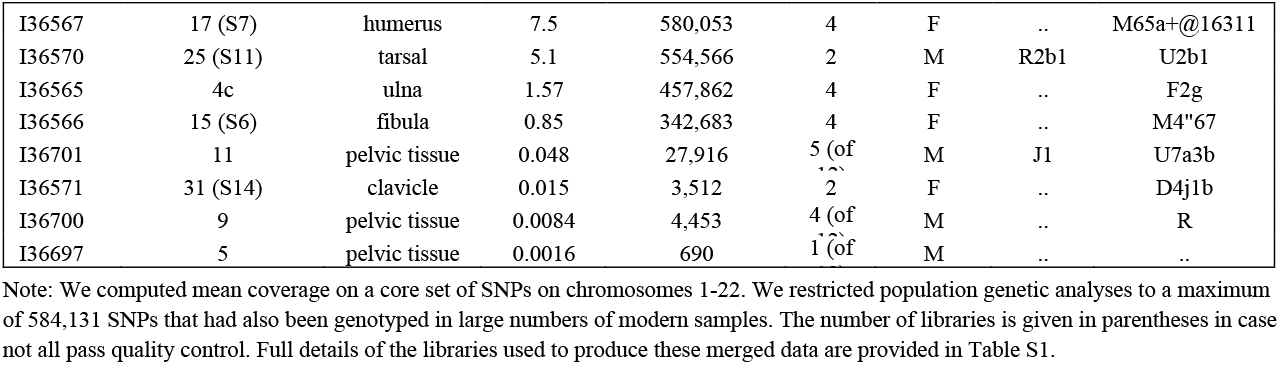
Summary of genetic results.

We determined genetic sex based on the ratio of Y to X chromosome sequences, identifying 6 genetic females and 4 genetic males (all three tissue samples were from males). Based on the ratio of the number of allelic differences between two sequences of the same individual, versus two sequences of different individuals (*18*), we determined that I36572 is a second degree relative of I36567, and a second-to-third degree relative of I36568.

We generated accelerator mass spectrometry (AMS) radiocarbon dates on the eight bone samples (Table S2). The 95% confidence intervals for the dates overlap in the range 1513-1347 calibrated years before present (*16*) (calBP). The union of 95% confidence intervals is 1690-1347 calBP.

### Ancestry intermediate between North Indians and Tibetans

We carried out Principal Component Analysis (PCA) (*19*) by co-analyzing genome-wide data from French, Han Chinese, several Tibetan plateau populations (modern Sherpa, modern Tibetans and 4600 year old Zongri (*20)*), and a variety of populations on the ‘Indian Cline’, referring to a previously described genetic gradient of present-day South Asian groups with relatively more (Pathan and North Indian Brahmins) and less (Palliyar) genetic sharing with West Eurasians (*21, 22*). We then projected the Old Lady Spider Cave individuals, who we call *Old Ladakh* in what follows, as well as relevant modern populations including Minero from Ladakh (*23*), Balti from Ladakh (*24*), and more than a thousand other individuals from diverse Indian populations that make up the Indian Cline including Dogra from Jammu and Kashmir (*21, 22*) (Table S3 gives the coordinates of each plotted point as well as their population affiliations). The first eigenvector separates people with primarily East Asian ancestry, and the second separates groups of the Indian Cline (Figure 1). A homogeneous cluster of the seven *Old Ladakh* individuals with highest quality data, as well as the Minero and Balti populations from the Himalayan region, fall along a line joining Tibetans to a point on the Indian Cline that includes both Dogra and North Indian Brahmins. This is the pattern expected if their ancestry is admixed between groups related to these populations. Except Balti and Minero, we are not aware of other modern groups plotting in a position similar to *Old Ladakh*, who have a unique ancestry profile for South Asia (**Figure 2**).

**Figure 2:**
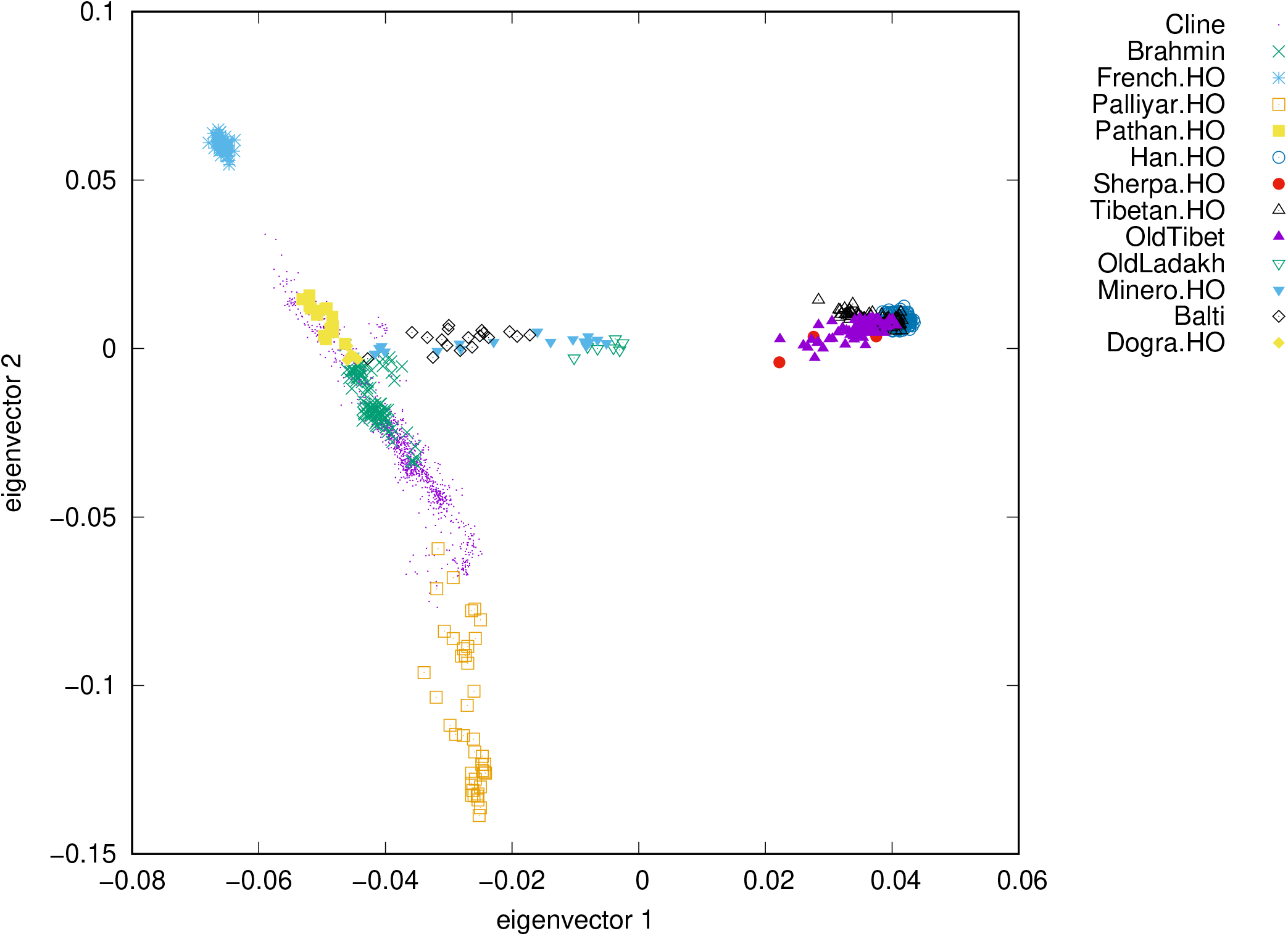
Principal Component Analysis reveals the relationship of *Old Ladakh* individuals to selected ancient and modern individuals. We define the axes of the PCA using selected Tibetan individuals from before the Iron Age (“OldTibet”), modern European and East Asian populations, and a variety of groups on the Indian Cline including Pathan (extreme northern end of the Cline), several North Brahmin groups, and Palliyar (at the extreme southern end). We project the seven higher coverage Old Ladakh individuals, as well as Dogra, Minero, Balti and 1153 individuals from diverse present-day Indian populations (tiny purple dots).

### No fitting two-source model for *Old Ladakh*

To quantify the mixture proportions and distinguish models, we used *qpAdm*, part of the ADMIXTOOLS software package (*17*). *qpAdm* allows us to propose a model for the pool of *Old Ladakh* individuals as a mixture of populations that can be proxied by a set of ‘left source’ populations hypothesized to be descended without mixture from the same origin as the true source populations. *qpAdm* also provides a formal statistical test of fit of the model to data. For our analyses, we often use modern populations as proxies for the ancient ones; as an example, motivated by the PCA, we used North Indian Brahmins as a source in our modeling of *Old Ladakh*, even though we do not think that Indians admixing into Ladakh were culturally ‘Brahmins’ and identifying these ancient people with modern cultural groups makes little sense. Using modern proxies as a source in *qpAdm* is well-motivated from a statistical point of view, however, and is a case for which *qpAdm* has a clear interpretation: the methodology was developed a decade ago for the application of studying European population history at a time when ancient DNA from that region were scarce as remains the case for South Asia today (*25*). If *qpAdm* produces a passing p-value, the proposed proxies for the sources are consistent with descending without mixture from the same groups as the true source populations. Even if the proxies postdate the true sources, we can use them to obtain unbiased estimates of mixture with valid standard errors.

To provide statistical leverage to falsify proposed models, *qpAdm* requires a set of ‘right outgroup’ populations related differentially to the modeled population and the left sources. For this application, we used eight right outgroups (Table S4). Four are modern groups genotyped on the Affymetrix Human Origins (HO) SNP array: 61 ‘French.HO’ individuals to represent Europeans (*17, 26, 27*); 107 ‘Han.HO’ individuals to represent East Asians (*17, 28);* 41 ‘Palliyar.HO’ individuals to represent a South Asian group on the Indian Cline with minimal West Eurasian relatedness (*22*), and 26 ‘Juang.HO’ individuals to represent a South Asian Austroasiatic-speaking group with even less West Eurasian relatedness (*22*). Four right outgroups are ancient: a pool of 10 diverse ancient Africans with little or no West Eurasian ancestry and dating to 8000-1000 BP ‘OldAfrica’ (*29-32);* 9 pastoralists from the western Zagros mountains site of Ganj Dareh dating to around 10000 BP ‘Iran_GanjDareh_N.AG’ (*27, 33, 34);* 21 hunter-gatherers dating to around 7000 BP from the middle to lower Volga region of Russia from the Ekaterinovka Mys site ‘Russia_Ekaterinovka_Eneolithic.AG’ (*34*); and 3 Mesolithic hunter-gatherers from the Amur River region of far northeastern China dating to around 11000 BP ‘China_AmurRiver_Mesolithic.AG’ (*35*).

We performed an initial analysis in which we tested 220 possible two-way mixture models for *Old Ladakh*, motivated by the position of these individuals in the PCA These models included as proxy for the ancient South Asian-related source one of the 4 North Brahmin groups, and, as a proxy for the Tibetan-related source, one of 55 modern or ancient East Asians groups (Table S4).

Table 2 shows all two source models for *Old Ladakh* fitting at P>0.001 where both have inferred ancestry proportions between 0-100% (Table S5 gives results for all models tested). No models are excellent, but the better ones all include one source related to Brahmins in North India contributing about 50% ancestry, and a second related to ancient Tibetans. The exception is that in the four best-fitting models, the non-South Asian source is an individual from the Amur River region of far northeastern China dating to ∼1650 BP (*36*). That we only observe good fits (P>0.05) using this Amur River individual should not be interpreted as strong evidence that the true source is from this region, however, since with only around 40,000 SNPs and a single individual, there is limited power to reject incorrect models. Since geographically proximate Tibetan populations fit second-best, but not fit exactly, we hypothesized that the true source is a Tibetan group that is not perfectly proxied by any populations we had available, perhaps one with more Amur River relatedness. Indeed, the degree of Amur River relatedness varied in ancient Tibetans: the statistic f_4_(OldAfrica, China_Amur-River_Mesolithic; Agangrong, Zhangcun) is significantly non-zero for the two ancient Tibetan groups Agangrong and Zhangcun (Z = -6.4). Similarly, previous work has shown that Tibetan groups varied in their degree of relatedness to ancient people from Xinjiang like Bronze Age Xiaohe (*20*). Based on these considerations, the most plausible explanation for our observations is that the true Tibetan source population had a slightly different mixture of ancestries than any of the ancient Tibetans for which we had available, but fell within their sampled variation.

**Table 2:**
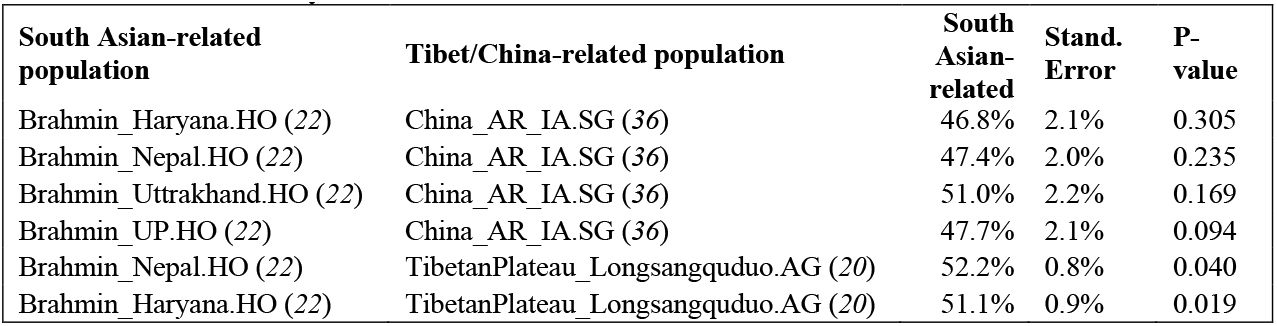
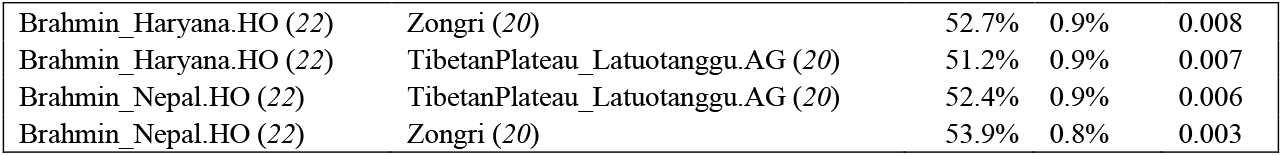
Best two-way mixture models for *Old Ladakh*. Full results in Table S5.

| South Asian-related population | Tibet/China-related population | South Asian-related | Stand. Error | P-value |
| --- | --- | --- | --- | --- |
| Brahmin_Haryana.HO (22) | China_AR_IA.SG (36) | 46.8% | 2.1% | 0.305 |
| Brahmin_Nepal.HO (22) | China_AR_IA.SG (36) | 47.4% | 2.0% | 0.235 |
| Brahmin_Uttarakhand.HO (22) | China_AR_IA.SG (36) | 51.0% | 2.2% | 0.169 |
| Brahmin_UP.HO (22) | China_AR_IA.SG (36) | 47.7% | 2.1% | 0.094 |
| Brahmin_Nepal.HO (22) | TibetanPlateau_Longsangquduo.AG (20) | 52.2% | 0.8% | 0.040 |
| Brahmin_Haryana.HO (22) | TibetanPlateau_Longsangquduo.AG (20) | 51.1% | 0.9% | 0.019 |
| Brahmin_Haryana.HO (22) | Zongri (20) | 52.7% | 0.9% | 0.008 |
| Brahmin_Haryana.HO (22) | TibetanPlateau_Latuotanggu.AG (20) | 51.2% | 0.9% | 0.007 |
| Brahmin_Nepal.HO (22) | TibetanPlateau_Latuotanggu.AG (20) | 52.4% | 0.9% | 0.006 |
| Brahmin_Nepal.HO (22) | Zongri (20) | 53.9% | 0.8% | 0.003 |

### *Old Ladakh* people had roughly half their ancestry from an unsampled population genetically similar to but not exactly the same as sampled ancient Tibetans

To identify fitting models for the ancestry of the *Old Ladakh* individuals, we tested 4940 3-source *qpAdm* models, considering all possible combinations of 260 South Asian diverse modern and ancient groups, 19 Tibetan modern and ancient groups, and people from the genetically distinctive Xiaohe culture from Bronze Age Xinjiang who lived approximately 5000-4800 BP (China_Xinjiang_Xiaohe_BA.AG) (Table S6) (*37*). For these analyses, we modified the right outgroup set used for two-source qpAdm models, removing Palliyar.HO, and adding three groups we found after exploration to provide additional leverage to distinguish between East Asian-related ancestry. These were a pool of 4 Late Paleolithic hunter-gatherers from the Amur River region of far northeastern China dating to 15000 BP ‘China_AmurRiver_LPaleolithic.AG’ (*35*); the low coverage Amur River region Iron Age individual dating to around 1800 BP who was part of the only fitting models in the 2-way *qpAdm* analysis (*36);* and a pool of 8 individuals from the Xiongnu culture dating to around 2000 BP ‘Russia_Buryatia_Xiongnu.AG’ (52).

We identified 80 three-way models fitting at P>0.05 that inferred mixture proportions for all three sources between 0-100% (results for all models including standard errors are in Table S7). Restricting to groups represented in more than one fitting model, all included a North Indian-related source (typically 46-50%), a Tibetan-related source (43-52%), and Xiaohe-related source (5-8%) (Table 3). The exceptions were Bahun.HO from Nepal (3 model fits) and Minero.HO from Ladakh (11 models fits), whose estimated contributions to fitting models were higher than for other South Asians at 59-65%, reflecting the fact they themselves are mixes of typically ‘Indian Cline’ and East Asian ancestry (22, 26, 43).

**Table 3:**
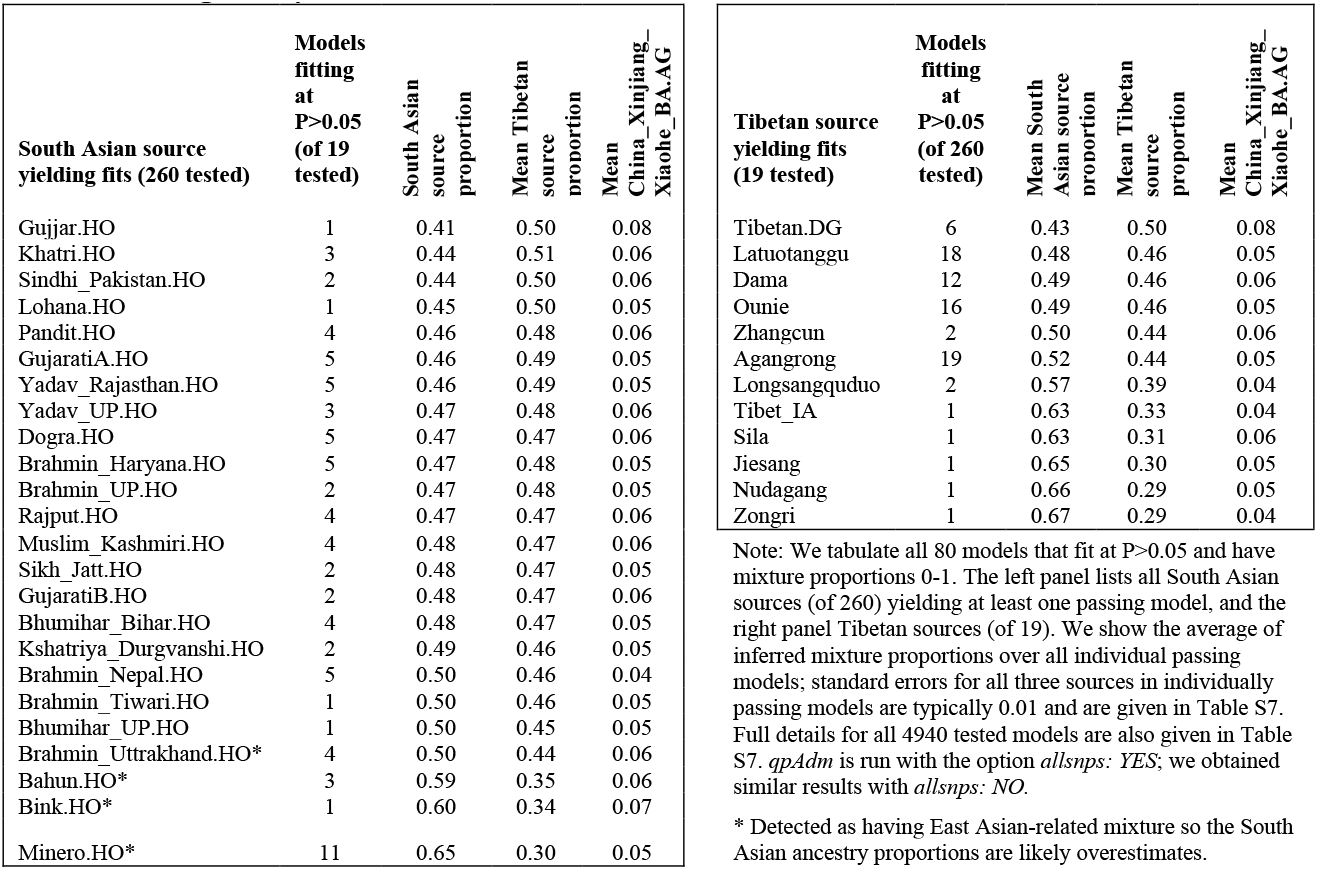
Fitting 3-way mixture models for *Old Ladakh*.

To probe the robustness of our best-fitting models, we focused on one that included North Indian Brahmins from Uttrakhand, ancient Tibetans from Latuotanggu, and ancient Xiaohe (P=0.17; Table S7). We then used the same eight right outgroups as for our primary *qpAdm* analysis, and carried out 492 *qpAdm* runs in turn, adding diverse populations to the Right. Table S8 shows the cases where moving a population to the right causes P-values that are significant after Bonferroni correction for the number of hypotheses tested. We observe some poor fits when adding to the right modern Himalayan populations such as Burusho.HO, Kalash.HO, and Minero.HO. This implies genetic drift shared between these groups and the *Old Ladakh* individuals, as expected for flow of *Old Ladakh-related* people into the ancestors of these groups. Thus, the *Old Ladakh* population did not completely disappear, and instead left a legacy in present-day people.

It is tempting to interpret the inclusion of 5-8% Xiaohe ancestry in our fitting models as evidence that this isolated Bronze Age group contributed ancestry to *Old Ladakh* people. However, this interpretation is not warranted and in fact is brought into question by our analyses. When we move to the right of our *qpAdm* model Shamanka Eneolithic from Russia, or Khovsgol Late Bronze Age from Mongolia, the models fail (Table S7), as would be expected for these groups having received gene flow from a Xinjiang or Amur River related source that lived later than the very ancient and isolated Xiaohe. The conservative interpretation of our results is that we do not have access to data from exactly the right Tibetan-related population, but that the true Tibetan-associated source population had a different mixture of ancestry ingredients present in Tibetan-related populations (Tibetan-associated, Xinjiang-associated, and Amur River-associated) compared to the specific individuals for which we have ancient DNA.

Taken together, our data are consistent with *Old Ladakh* individuals being a mixture of approximately half ancestry related to North Brahmins and Dogra, and half ancestry related to sampled ancient Tibetans but not perfectly proxied by any available Tibetans in our dataset. These results are also compatible with the patterns observed at uniparentally transmitted mitochondrial DNA sequences and Y chromosomes (Table 1). Six haplogroups are typically South Asian and nearly absent in Tibetans: both the Y haplogroups R2b1 and J1, and 4 mitochondrial haplogroups M65a+@16311, U2b1, M4”67 U7a3b (https://www.yfull.com/tree/). Two mitochondrial haplogroups are common in Tibetans, and nearly absent in South Asians: F2g and D4j1b. One mitochondrial haplogroup, R, is consistent with either a South Asian or a Tibetan source. Finally, a pair of 2^nd^ to 3^rd^ degree relatives carry mitochondrial haplogroup U5a1d1, characteristic of modern and ancient North Europeans including from the Caucasus and Lower Volga area, but both are rare in South Asia today. The most plausible hypothesis is that this haplogroup arrived in North India as a low frequency allele carried by people with steppe ancestry.

### *Old Ladakh* ancestors formed through mixture by around 2800 BP

To estimate when the mixture of South Asian- and Tibetan-related ancestry occurred, we ran the software DATES (*38, 39*). We used a pool of 35 North Indian Brahmin individuals from four areas (Haryana, Nepal, Tiwari, Uttrakhand, Uttar Pradesh) as a proxy source to represent ancient South Asians, and 143 modern people with Tibetan-related ancestry as a proxy source to represent ancient Tibetans. DATES reveals a clear covariance of ancestry with distance (**Figure 3**). The data are well fit by a single exponential function, implying the population was insulated from later waves of mixture. Assuming 29 years per generation, we infer the admixture occurred 1343±140 years before the date of the individuals, or roughly 2800 BP. Previous work showed that Iron Age people from what is now northern Pakistan had all three of the deep ancestry components that contributed to the modern Indian Cline by this time, including Steppe-related, Iran-related, and indigenous Ancient South Indian-related (*38*). However, those data did not have these ancestries in the right proportions to be on the Indian Cline. The ancestry of the *Old Ladakh* individuals shows the Indian Cline—people with ancestry at the location of the Indian Cline where present-day Dogra and North Indian Brahmins fall—was in place by ∼2800 BP.

**Figure 3:**
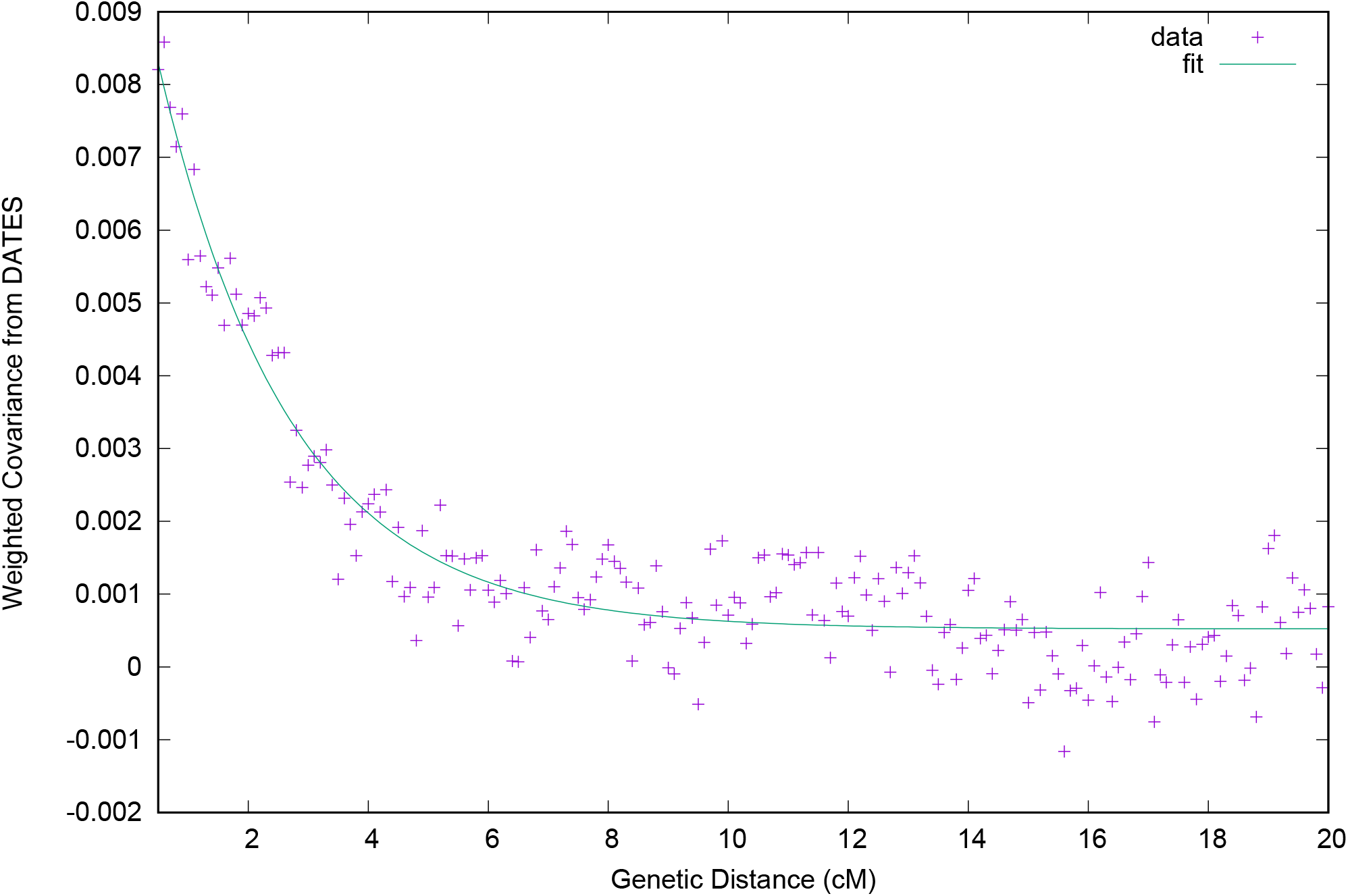
Correlation of ancestry with genetic distance in the *Old Ladakh* individuals, inferred by. We observe a decay of admixture linkage disequilibrium on the scale of 2 centimorgans, implying that admixture began at least 50 generations before they lived.

### *Old Ladakh* people carried a variant from Denisovans conferring high altitude adaptation

Modern and ancient Tibetans frequently carry an allele that confers adaptation to a high-altitude environment, which has been shown to have introgressed from archaic Denisovans (*40*). To test for the presence of the Denisovan haplotype carrying this allele in the newly sequenced individuals, we used the raw data from the individuals we sequenced to identify 11 SNPs that are diagnostic for the Denisovan haplotype and are effectively ascertained after our enrichment process. One individual has no coverage on these SNPs so we can make no call. Another has just a single allele (not on the haplotype) so we make a call on just 9 chromosomes from the (diploid) individuals from Old Ladakh. Of these, 1 carries the Denisovan haplotype. Approximately half of the Ancient Tibetan chromosomes have this haplotype and we estimate that Ancient Tibetans contributed about half the ancestry of the *Old Ladakh* individuals. Thus, in the absence of natural selection, about a quarter of chromosomes of Old Ladakh are expected to have the Denisovan haplotype, which gives an expected number as 2.25. Seeing only 1 is not statistically surprising. All we can say is that after admixture this haplotype was not under strong positive selection.

## Discussion

The ancient individuals we analyzed from Ladakh are of great interest. First, there is little genetic data available from anywhere in the Indian subcontinent that is as old as these individuals. Second, their ancestry is unusual: very few groups today have a similar ∼50-50% South Asian/Tibetan mix. Third, we are likely learning about the deep history of Ladakh, although a caveat is that the very old admixture did not necessarily occur in Ladakh itself. For example, admixture could have occurred in what is now Nepal, and then members of the admixed population moved to Ladakh. Fourth, the data provide direct evidence of contacts between Ladakh and Tibet a century before the rise of the Tibetan Empire (*41*).

Our results do not imply that the detected admixture took place in Ladakh. However, we have shown that the people from Old Lady Spider Cave were part of a population that was long-established—arising through admixture at least around 1300 years before the time the sample people lived—and likely made a contribution to people living today in the Himalayan region including Ladakh. In particular, groups like the Minero from Ladakh have *qpAdm-based* evidence of gene from an *Old Ladakh* group (Table S8), and they and Balti fall on a gradient of differential relatedness between North Indian groups and *Old Ladakh* (Figure 2). In our *qpAdm* analysis, Minero fit well (P=0.24) as a mixture of Dogra (44±1%) and *Old Ladakh* (56±1%), and Balti fit marginally (P=0.04) as a mixture of Dogra (55±1%) and *Old Ladakh* (45±1%). An important direction for future work is to sample additional remains to understand the geographic and temporal span, as well as the cultural associations, of this previously unknown population that lived in the Himalayan region for more than a millennium.

## Materials and Methods

### Provenance

Excavation of the human remains was carried out in 2021 by a team led by two of the corresponding authors (Veena Mushrif-Tripathy and Quentin Devers), under the auspices of the Archaeological Survey of India (ASI), which gave permission for this work (*9*). Permission for sending the archaeological materials to Harvard University for specialized ancient DNA analysis and radiocarbon dating was obtained from the Archaeological Survey of India in a letter dated December 30 2021, and they were hand-carried by corresponding author Veena Musrif-Tripathy. Open science principles require making all data used to support the conclusions of a study maximally available, and we support these principles here by making fully publicly available not only the digital copies of molecules (the uploaded sequences) but also the molecular copies (the ancient DNA libraries themselves, which constitute molecular data storage, and reside at Harvard Medical School). Those researchers who wish to carry out deeper sequencing of libraries published in this study should make a request to corresponding author D.R. We commit to granting reasonable requests as long as the libraries remain preserved in our laboratories, with no requirement that we be included as collaborators or co-authors on any resulting publications. The remaining skeletal samples are housed at Deccan College, and we commit to granting reasonable requests to study them from an anthropological point of view; all queries related to anthropological study should be made to corresponding author V.M.-T.

### Ancient DNA wet laboratory work

The ancient DNA generation methods we used are referenced in the main text and specified on a per-library basis in Table S1. For the specialized extraction procedures we used for the soft tissue remains, we tried four different extraction buffers for each sample:

Buffer 1: 0.45M EDTA (pH 8.0), 0.05% Tween-20 and 0.25mg/ml proteinase K (*10*).

Buffer 2: 4.2M guanidine isothiocyanate, 0.053 M Tris-HCl (pH 7.5), 0.0106 M EDTA, and 2.12% Sarkosyl and 0.2mg/ml proteinase K (*11*).

Buffer 3: 0.01M Tris-HCl (pH 8.0), 0.01M NaCl, 5mM CaCl2, 2.5mM EDTA (pH 8.0), 2% w/v SDS, 0.04 M Dithiothreitol (DTT) and 10% proteinase K solution (>600 mAU/ml, Qiagen)(*12*)

Buffer 4: ATL buffer and 3% proteinase K solution, both from the DNeasy Blood and Tissue Kit (Qiagen) (*13*).

Proteinase K was added just before adding buffer to samples. 37 mg of tissue was cut into small fragments (about 5 cubic mm) and homogenized in 750μl of the selected extraction buffer using a pestle in a 2 ml Eppendorf low-bind DNA tube. The samples were incubated overnight at 56°C (rotated). DNA extraction followed previously published protocols for different lysis buffers (*10-13*), with one modification: silica beads were used for purification instead of columns. Doublestranded libraries were prepared from the DNA extracts following an established protocol (*14*). To increase sequence coverage, single-stranded libraries were generated for most extracts, including from bone (*15, 42*). To reduce damage-induced errors, libraries were treated with uracil-DNA glycosylase (UDG) to cleave ancient DNA molecules at uracils on the 5′ end, a damage modality characteristic of ancient DNA (*14, 43*). Libraries were then enriched for sequences overlapping the mitochondrial genome and approximately 1.4 million genome-wide SNPs (*16*). Double-stranded libraries underwent one round of capture, and single-stranded libraries two rounds, using the reagents and buffers produced by Twist Biosciences (*16*). Successful double-strand libraries were sequenced on the Illumina NovaSeq S4 platform with paired-end reads, while single-strand libraries were sequenced on the Illumina NovaSeq X platform, also with paired-end reads.

### Radiocarbon Dating

We obtained 8 accelerator mass spectrometry radiocarbon dates for 7 distinct individuals from the Pennsylvania State University Radiocarbon Laboratory. To generate these dates, we sonicated bone samples in successive washes of American Chemical Society-grade methanol, acetone and dichloromethane for 30 min each at room temperature and followed this by three washes in Nanopure water to remove possible contaminants (including conservants and adhesives). We extracted bone collagen and purified it using a modified Longin method with ultrafiltration (>30 kDa gelatin) (*44*). We measured carbon and nitrogen concentrations and C/N ratios of the extracted and purified collagen samples using a Costech elemental analyzer (ECS 4010) and evaluated sample quality by percentage of crude gelatin yield, percentage of C, percentage of N, and C/N ratios before accelerator mass spectrometry 14C dating. C/N ratios for all samples fell between 3.1 and 3.3, indicating excellent preservation (*45*). We combusted collagen samples for 3 hours at 900 degrees C in vacuum-sealed quartz tubes with CuO and Ag wires. We reduced sample CO_2_ to graphite at 550 degrees C using H_2_ and a Fe catalyst, and we drew off reaction water with Mg(ClO_4_) (*46*).

We pressed graphite samples into targets in aluminum boats, loaded them onto a target wheel, and performed all measurements using a modified National Electronics Corporation compact spectrometer with a 0.5 MV accelerator (NEC 1.5SDH-1). We corrected 14C ages for massdependent fractionation with measured 513C values (*47*) and compared these with samples of whale bone from the Pleistocene (>48000 BP), bison bone from the late Holocene (around 1850 calibrated years BP), cow bone from the late 1800s CE, and OX-2 oxalic acid standards. We calibrated radiocarbon ages using OxCal v.4.45 and the IntCal20 Northern Hemisphere curve (*48, 49*). For one individual (I36568), we generated two overlapping dates using different bone elements, and used the RCombine function in OxCal to combine these into a single weighted average. We report the 95.4% Bayesian credible intervals for all dates (Table S2).

## Supporting information

Supplementary Tables 1-8 in Excel format

## SUPPLEMENTARY MATERIALS

Supplementary material for this article is available at [address to be made available upon publication].

## Acknowledgments

The research team in the field was composed of Veena Mushrif-Tripathy, Quentin Devers, Sonam Dolma, Sumeet Sunil, and Saurabh Singh. We are grateful for support from the beginning of the project from Kacho Mumtaz Khan, Kacho Sikundar Khan, and Kacho Shamim Khan, and also to Dr. Kacho Akbar Khan, Kacho Asfandyar Khan, Tashi Ldawa Tshangspa, Sultan Wazir, Viraf Mehta, and Sonam Choldan (Gasha-pa). We are grateful for the administrative support of Kacho Mehboob Khan (Cultural Secretary, UT Ladakh), Dr. Shivdardhan Singh Jamwal (Additional Director General of Police, UT Jammu & Kashmir), Kacho Ishtyaq Khan (Deputy Superintendent of Police, Leh), Santosh Sukhadeve (District Commissioner, Kargil), Shrikant Balasaheb Suse (District Commissioner, Leh), and Avny Lavasa (previous District Commissioner, Leh). We are thankful for the help in the field of Ibrahim Mohammad (Pata-pa), Sadiq Ali (Pata-pa), Ashiq Hussain (Shanak-pa), Ali Sher (Khan-pa), Jamshed Hussain (Pata-pa), Marzia Bano (Dengo-ma), Mohammad Amin (Dengo-pa), Sogra Fatima (Dengo-ma), Mohammad Mussa (Ayid-pa). We appreciate discussions with Samara Broglia de Moura, Nicolas Tournadre, Guillaume Jacques, Alexandre Beylier, and Pauline Sebillaud. We are grateful to Lomous Kumar and Niraj Rai for sharing the Minero data from their paper. We thank Daniel Tabin for advice on Turkic and Mongolian history and suggestions for populations for co-analyzing Amur River groups and Tibet. We thank Kendra Sirak for help in interpreting radiocarbon dates.

## Author contributions

Conceptualization: N.P., V.M.-T., Q.D., and D.R. Supervision: V. M.-T., S.M., N.R., and D.R. Archaeological material: V.M.-T., Q.D., and S.D. Contextualization: V.M.-T. and Q.D. Visualization: N.P., V.M.-T., Q.D., and D.R. Writing: N.P., V.M.-T., Q.D., and D.R.

## Data and materials availability

All data needed to evaluate the conclusions in the paper are present in the paper and/or the Supplementary Materials. Newly reported ancient sequencing data in this study will be deposited at the European Nucleotide Archive (ENA) and will be publicly available as of the date of publication. The accepted version of this article (before the editing, proofreading and formatting changes following the paper being accepted) is subject to the Howard Hughes Medical Institute (HHMI) Open Access to Publications policy; HHMI lab heads have previously granted a non-exclusive CC BY 4.0 license to the public and a sublicensable license to HHMI in their research articles. Pursuant to those licenses, the accepted manuscript can be made freely available under a CC BY 4.0 license immediately on publication.

## Supplementary Table Captions

**Table S1:** Details of DNA extraction, library preparation, and sequencing for 64 newly generated libraries

**Table S2:** Details of 10 newly generated radiocarbon dates and associated isotopic measurements

**Table S3:** Details of 1696 individuals used in Principal Component Analysis (Figure 2)

**Table S4:** Details of 1094 individuals used in the 2-source qpAdm analysis

**Table S5:** Full results for 2-source qpAdm models

**Table S6:** Details of 2685 individuals used in the 3-source qpAdm analysis

**Table S7:** Full results for 3-source qpAdm models

**Table S8:** Addition of 492 populations in turn to the right outgroups for the Brahmins_Uttrakhand.HO-Latuotanggu-China_Xinjiang_Xiaohe_BA.AG qpAdm model to probe model robustness

## REFERENCES AND NOTES

1. H.-P. K. Francfort, Daniel; Mascle, George, in Rock art in the old world: papers presented in Symposium A of the AURA Congress, Darwin (Australia), 1988, M. Lorblanchet, Ed. (Indira Gandhi National Centre for the Arts : UBS Publishers’ Distributors IGNCA rock art series, New Delhi, 1992), pp. 147–192.

2. L. Bruneau, Universitě Paris 1 Panthéon-Sorbonne, (2010).

3. L. Bruneau, The Rock Art of Ladakh: A Historiographic and Thematic Study. Rock Art: Recent Researches and New Perspectives (Festschrift to Padma Shri Dr. Prof. Yashodhar Mathpal), 79–99 (2015).

4. S. Broglia de Moura, Preliminary field notes on the ancient ceramics of Ladakh. Etudes Mongoles et Siberiennes 51, (2020).

5. S. Broglia de Moura, Universitě PSL, (2022).

6. Archaeological Ladakh: Recent Discoveries Redefining the History of a Key Region between the Pamirs and the Himalayas. Central Asiatic Journal 61, (2018).

7. Q. Devers, Historical Sites of Nubra, Leh, Ladakh. (Indian National Trust for Art and Cultural Heritage, Delhi, 2021).

8. R. Salomon, Q. Devers, T. Ldawa, Kharosthi and Brähmi Inscriptions from Ladakh. Bulletin of the Asia institute / Asia institute (Detroit, Mich.), (2021).

9. V. Mushrif-Tripathy, Q. Devers, S. Dolma, K. Mumtaz Khan, K. Sikundar Khan, The funerary cave of the Old Lady Spider (Ladakh). Revue d’Étude Tibétaine, Forthcoming (2026).

10. N. Rohland, I. Glocke, A. Aximu-Petri, M. Meyer, Extraction of highly degraded DNA from ancient bones, teeth and sediments for high-throughput sequencing. Nat Protoc 13, 2447–2461 (2018).

11. M. M. Tin, E. P. Economo, A. S. Mikheyev, Sequencing degraded DNA from non-destructively sampled museum specimens for RAD-tagging and low-coverage shotgun phylogenetics. PLoS One 9, e96793 (2014).

12. P. Campos, T. Gilbert, DNA Extraction from Keratin and Chitin. Methods in molecular biology (Clifton, N. J.) 840, 43–49 (2012).

13. A. Gómez-Carballa et al., The complete mitogenome of a 500-year-old Inca child mummy. Scientific Reports 5, 16462 (2015).

14. N. Rohland, E. Harney, S. Mallick, S. Nordenfelt, D. Reich, Partial uracil-DNA-glycosylase treatment for screening of ancient DNA. Philosophical transactions of the Royal Society of London. Series B, Biological sciences 370, 20130624 (2015).

15. M.-T. Gansauge, A. Aximu-Petri, S. Nagel, M. Meyer, Manual and automated preparation of single-stranded DNA libraries for the sequencing of DNA from ancient biological remains and other sources of highly degraded DNA. Nature Protocols 15, 2279–2300 (2020).

16. N. Rohland et al., Three assays for in-solution enrichment of ancient human DNA at more than a million SNPs. Genome Res 32, 2068–2078 (2022).

17. N. Patterson et al., Ancient admixture in human history. Genetics 192, 1065–1093 (2012).

18. D. J. Kennett et al., Archaeogenomic evidence reveals prehistoric matrilineal dynasty. Nat Commun 8, 14115 (2017).

19. N. Patterson, A. L. Price, D. Reich, Population structure and eigenanalysis. PLoS genetics 2, e190 (2006).

20. H. Wang et al., Human genetic history on the Tibetan Plateau in the past 5100 years. Sci Adv 9, eadd5582 (2023).

21. D. Reich, K. Thangaraj, N. Patterson, A. L. Price, L. Singh, Reconstructing Indian population history. Nature 461, 489–494 (2009).

22. N. Nakatsuka et al., The promise of discovering population-specific disease-associated genes in South Asia. Nat. Genet. 49, 1403–1407 (2017).

23. L. Kumar et al., The genetic landscape, gene flow and adaptation in Ladakh highlanders of trans-Himalayas. bioRxiv, 2024.2002.2005.579041 (2024).

24. X.-Y. Yang et al., Tracing the Genetic Legacy of the Tibetan Empire in the Balti. Molecular Biology and Evolution 38, 1529–1536 (2021).

25. W. Haak et al., Massive migration from the steppe was a source for Indo-European languages in Europe. Nature 522, 207–211 (2015).

26. I. Lazaridis et al., Ancient human genomes suggest three ancestral populations for present-day Europeans. Nature 513, 409–413 (2014).

27. I. Lazaridis et al., Genomic insights into the origin of farming in the ancient Near East. Nature 536, 419–424 (2016).

28. C. C. Wang et al., Genomic insights into the formation of human populations in East Asia. Nature 591, 413–419 (2021).

29. M. Lipson et al., Ancient West African foragers in the context of African population history. Nature 577, 665–670 (2020).

30. P. Skoglund et al., Reconstructing Prehistoric African Population Structure. Cell 171, 59-71.e21 (2017).

31. K. Wang et al., Ancient genomes reveal complex patterns of population movement, interaction, and replacement in sub-Saharan Africa. Sci Adv 6, eaaz0183 (2020).

32. M. Lipson et al., Ancient DNA and deep population structure in sub-Saharan African foragers. Nature 603, 290–296 (2022).

33. V. M. Narasimhan et al., The Genomic Formation of South and Central Asia. bioRxiv, (2018).

34. I. Lazaridis et al., The genetic origin of the Indo-Europeans. Nature 639, 132–142 (2025).

35. X. Mao et al., The deep population history of northern East Asia from the Late Pleistocene to the Holocene. Cell 184, 3256-3266.e3213 (2021).

36. C. Ning et al., Ancient genomes from northern China suggest links between subsistence changes and human migration. Nat Commun 11, 2700 (2020).

37. F. Zhang et al., The genomic origins of the Bronze Age Tarim Basin mummies. Nature 599, 256–261 (2021).

38. V. M. Narasimhan et al., The formation of human populations in South and Central Asia. Science 365, eaat7487 (2019).

39. M. Chintalapati, N. Patterson, P. Mooijani, The spatiotemporal patterns of major human admixture events during the European Holocene. eLife 11, e77625 (2022).

40. E. Huerta-Sanchez et al., Altitude adaptation in Tibetans caused by introgression of Denisovan-like DNA. Nature 512, 194–197 (2014).

41. C. I. Beckwith, The Tibetan empire in central Asia : a history of the struggle for great power among Tibetans, Turks, Arabs, and Chinese during the early Middle Ages. (Princeton University Press, Princeton, N.J., 1993), pp. xxii, 281 p.

42. M. T. Gansauge et al., Single-stranded DNA library preparation from highly degraded DNA using T4 DNA ligase. Nucleic Acids Res 45, e79 (2017).

43. A. W. Briggs et al., Removal of deaminated cytosines and detection of in vivo methylation in ancient DNA. Nucleic Acids Res 38, e87 (2010).

44. S. B. McClure, O. G. Puchol, B. J. Culleton, Ams Dating of Human Bone from Cova De La Pastora: New Evidence of Ritual Continuity in the Prehistory of Eastern Spain. Radiocarbon 52, 25–32 (2010).

45. G. J. Van Klinken, Bone collagen quality indicators for palaeodietary and radiocarbon measurements. J. Archaeol. Sci. 26, 687–695 (1999).

46. G. M. Santos, J. R. Southon, K. C. Druffel-Rodriguez, S. Griffin, M. Mazon, Magnesium Perchlorate as an Alternative Water Trap in AMS Graphite Sample Preparation: A Report On Sample Preparation at Kccams at the University of California, Irvine. Radiocarbon 46, 165–173 (2004).

47. M. Stuiver, Polach, H. A., Reporting of C-14 Data-Discussion. Radiocarbon 19, 355–363 (1977).

48. C. B. Ramsey, S. Lee, Recent and Planned Developments of the Program OxCal. Radiocarbon 55, 720–730 (2013).

49. P. J. Reimer et al., The IntCal20 Northern Hemisphere Radiocarbon Age Calibration Curve (0-55 cal kBP). Radiocarbon 62, 725–757 (2020).

52. C. Jeong, K. Wang, S. Wilkin, W. T. T. Taylor, B. K. Miller, J. H. Bemmann, R. Stahl, C. Chiovelli, F. Knolle, S. Ulziibayar, D. Khatanbaatar, D. Erdenebaatar, U. Erdenebat, A. Ochir, G. Ankhsanaa, C. Vanchigdash, B. Ochir, C. Munkhbayar, D. Tumen, A. Kovalev, N. Kradin, B. A. Bazarov, D. A. Miyagashev, P. B. Konovalov, E. Zhambaltarova, A. V. Miller, W. Haak, S. Schiffels, J. Krause, N. Boivin, M. Erdene, J. Hendy, C. Warinner, A Dynamic 6,000-Year Genetic History of Eurasia’s Eastern Steppe. Cell, 2020 183, 890–904.

